# Approach-Avoidance Bias in Virtual and Real-World Simulations: Insights from a Systematic Review of Experimental Setups

**DOI:** 10.1101/2024.12.12.628144

**Authors:** Aitana Grasso-Cladera, John Madrid-Carvajal, Sven Walter, Peter König

**Affiliations:** Institut für Kognitionswissenschaft. Osnabrück University. Osnabrück (49090), Germany; Department of Neurophysiology and Pathophysiology, Center of Experimental Medicine, University Medical Center Hamburg-Eppendorf, Hamburg (20251), Germany

**Keywords:** Approach-Avoidance Bias, Virtual Reality, Ecological Validity, Systematic Review, Embodied Cognition

## Abstract

**Background:** Approach and avoidance bias (AAB) describes automatic behavioral tendencies to react toward environmental stimuli regarding their emotional valence. Traditional setups have provided evidence but often lack ecological validity. The study of the AAB in naturalistic contexts has recently increased, revealing significant methodological challenges. This systematic review evaluates the use of virtual reality (VR) and real-world setups to study the AAB, summarizing methodological innovations and challenges.

**Methods:** We systematically reviewed peer-reviewed articles employing VR and real-world setups to investigate the AAB. We analyzed experimental designs, stimuli, response metrics, and technical aspects to assess their alignment with research objectives and identify limitations.

**Results:** This review included 21 studies revealing diverse methodologies, stimulus types, and novel behavioral responses, highlighting significant variability in design strategies and methodological coherence. Several studies used traditional reaction time measures yet varied in their application of VR technology and participant interaction paradigms. Some studies showed discrepancies between simulated and natural bodily actions, while others showcased more integrated approaches that preserved their integrity. Only a minority of studies included control conditions or acquired (neuro)physiological data.

**Conclusions:** VR offers a potential ecological setup for studying the AAB, enabling dynamic and immersive interactions. Our results underscore the importance of establishing a coherent framework for investigating the AAB tendencies using VR. Addressing the foundational challenges of developing baseline principles that guide VR-based designs to study the AAB within naturalistic contexts is essential for advancing the AAB research and application. This will ultimately contribute to more reliable and reproducible experimental paradigms and develop effective interventions that help individuals recognize and change their biases, fostering more balanced behaviors.

## Introduction

Approach-avoidance behaviors are automatic responses that organisms display toward environmental stimuli, influenced by the positive or negative value associated with these stimuli [1,2]. Organisms continuously interact with their surroundings, assessing objects, events, and opportunities based on perceived benefits or threats. The inclination to approach or avoid is guided by the valence, or emotional value, of stimuli: organisms tend to approach stimuli that are seen as rewarding or beneficial for safety and well-being [3–5]. Conversely, organisms tend to avoid stimuli linked to adverse or harmful outcomes, aiming to protect themselves from threats that could compromise their safety [3,6]. Emotional valence is central to regulating approach-avoidance behaviors, with responses driven by an underlying positive-negative valence behavioral system that influences interactions in meaningful ways [1,7,8].

### Approach-Avoidance Bias (AAB)

Empirical evidence regarding approach-avoidance behaviors has revealed that these actions are susceptible to reaction times effect and that an embodied component might be associated with it. Studies indicate that approach-avoidance behaviors are susceptible to reaction times, with organisms responding faster and more accurately to stimuli aligned with their instincts—approaching positive stimuli and avoiding negative ones—than to stimuli that contradict these predispositions [9–14]. This Automatic Approach-Avoidance Bias (AAB) suggests an inherited tendency to perform actions automatically, guided by perceptual, experiential, or cognitive biases [6]. Recent research highlights an inherent embodied component of the AAB, rooted in its evolutionary role, emphasizing the need to study it in dynamic, realistic contexts [6,15]. The embodied aspect of the AAB suggests that physiological mechanisms regulate these automatic actions, offering insights into how bodily dynamics shape decision-making processes and responses to environmental stimuli.

Research on the AAB encompasses diverse methodological approaches, employing various stimuli, tasks, and measures to address a wide range of questions. A key method is the Stimulus-Response Compatibility (SRC) task, particularly the widely used Approach-Avoidance Task (AAT), where participants perform an action symbolizing a specific movement (e.g., pushing or pulling via button pressing) or actually performing the actions using, for example, a joystick, in response to emotionally valenced stimuli while their reaction times are measured. Standard behavioral metrics to measure an AAB include response time (i.e., time taken to perform a specified action) and accuracy (i.e., alignment of action with instruction) [5,11]. Additional measures, such as ocular dynamics (e.g., saccades, fixations) for attentional processes [16,17] and autonomic nervous system activity —cardiac activity [18] and skin conductance [19]— have provided insights into vagal tone, stress responses, and emotion regulation [20–22]. This diverse set of techniques and metrics enriches our understanding of the AAB by enabling the study of its behavioral and physiological underpinnings across varied contexts.

The AAT has been one of the preferred paradigms for studying the AAB due to its versatility in fitting different topics. Nevertheless, the AAT and its variations have primarily been implemented as stationary, computer-based tasks that are highly controlled and tend to present non-realistic scenarios with fewer degrees of ecological validity, i.e., low representativity of stimuli and relevance of tasks within the context of functional action [23]. In this sense, real-world paradigms can offer newer insights on embodied aspects of the AAB (i.e., different bodily actions for approaching or avoiding and physiological dynamics), which are fundamental to further understanding, for example, motivation processes and the evolutive role of the bias [9]. Overall, Virtual Reality (VR) and real-life setups offer researchers the opportunity to deviate from classical paradigms by creating realistic scenarios to study the AAB, under controlled yet ecologically valid conditions.

The VR setup enables researchers to collect detailed behavioral data, including head, eye, hand, and body movements when participants interact with virtual objects—providing insights beyond just reaction time and response accuracy with regard to the embodiment of the AAB [5,11]. Also, VR devices open up the possibility of being combined with other measurement devices, facilitating the acquisition of multimodal data, such as from physiology, e.g., electrical brain and cardiac activity [24–26]. By implementing experimental paradigms based on VR opens the door for new research questions, allowing a paradigm shift to further understand the behavioral and embodied mechanisms involved in the AAB by performing naturalistic actions in a scenario closer to real life.

Previous research has demonstrated the feasibility of bringing the AAB study into VR. For example, Degner and collaborators [1] assessed the AAB in an immersive virtual environment, where they could replicate the main findings reported for desktop-based paradigms for the AAB. Also, the use of VR in the context of the AAB has been reported for the development of innovative treatments for different conditions (e.g., nicotine addiction, and eating disorders). An example of this is the line of work conducted by Eiler and collaborators [4,27,28], who have systematically implemented an Approach-Avoidance paradigm in VR to generate a Cognitive Bias Modification (CBM; i.e., an approach to modify automatic cognitive processes in a specific direction [29]) paradigm for the treatment of smoke/nicotine addiction [30]. Hence, VR’s integration into AAB research confirms its feasibility for both replicating traditional findings and advancing therapeutic applications. These developments underscore VR’s potential as a valuable tool for treatments.

However, the implementation of the paradigms in VR is not trivial and requires the assessment of methodologies and the overcoming of several challenges. For instance, a preliminary evaluation of transferring the AAB research into VR focusing on improved usability, functionalities, and graphics showed that Leap Motion sensors are preferred to controllers [4]. This transfer brings challenges such as the accuracy of time recording, the gesture or action to be performed and recorded, as well as rethinking the inclusion of the analysis of eye, head, hand, and body movements in the light of the AAB. Hence, issues such as timing accuracy, choice of gestures, and the use of sensors like Leap Motion over controllers highlight the need for advanced usability, precise tracking, and comprehensive analysis of movements to fully adapt AAB studies to virtual settings.

Considering this, the present review provides a comprehensive overview of the current literature that employs VR or real-life setups to study the AAB. Specifically, we focused on summarizing and examining methodological aspects of the existing literature. This thorough examination of the methods employed seeks to provide insights into how studies have approached the transfer of the AAB into VR or real-life setups. Also, this review sought to highlight the major limitations and challenges associated with using VR to study the AAB. Overall, with this systematic review, we attempted to establish a path for future research studying the AAB in setups with higher ecological validity, for a proper understanding of the mechanisms underlying the AAB and its embodied dynamics.

## Methods

The present review was conducted following the Joanna Briggs Institute (JBI) guidelines for systematic reviews and meta-analysis [31] and is reported following the Preferred Reporting Items for Systematic Reviews and Meta-analyses (PRISMA) [32,33].

### Eligibility criteria

This review focuses on studies conducted either in VR environments or in natural settings with a high degree of ecological validity to explore the phenomenon of Approach-Avoidance Bias. Given the diverse range of experimental designs in neuroscience and cognitive science research, no specific inclusion or exclusion criteria were applied based on the study design. As a result, all experimental and quantitative study designs published in peer-reviewed journals were included. However, due to the focus on methodology and analysis, study protocols were excluded. In terms of sample characteristics, only studies involving human populations were considered. This includes research on both healthy individuals and specific target groups (e.g., those with clinical conditions).

Since this review emphasizes the methodological and analytical aspects of Approach-Avoidance Bias, studies using an Approach-Avoidance Task in Virtual Reality or natural settings as a training or treatment tool for specific conditions were excluded. All other experimental paradigms aimed at studying Approach-Avoidance Bias in these settings were eligible for inclusion. Additionally, there were no restrictions related to comparison conditions, meaning the review includes studies with or without control conditions.

There were no time restrictions, and only studies published in English were considered. Table 1 provides a summary of the study selection criteria.

**Table 1:**
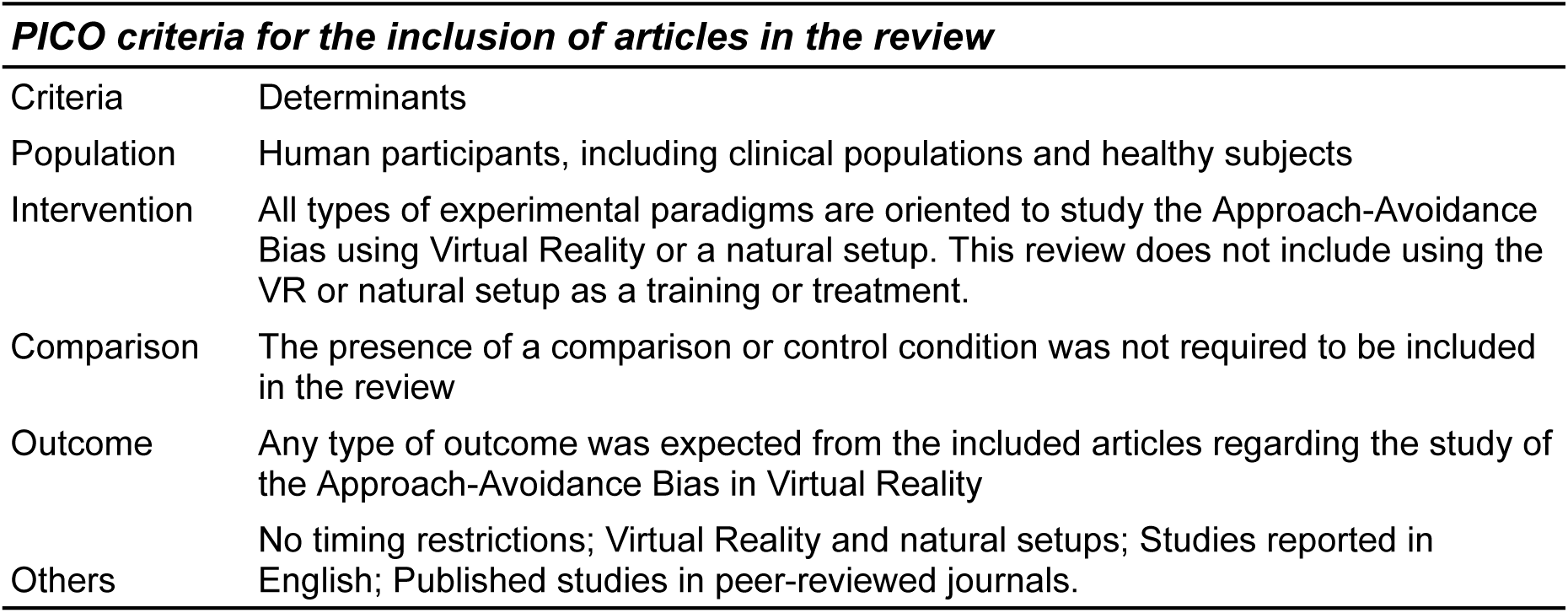

| ***PICO criteria for the inclusion of articles in the review*** |  |
| --- | --- |
| Criteria | Determinants |
| Population | Human participants, including clinical populations and healthy subjects |
| Intervention | All types of experimental paradigms are oriented to study the Approach-Avoidance Bias using Virtual Reality or a natural setup. This review does not include using the VR or natural setup as a training or treatment. |
| Comparison | The presence of a comparison or control condition was not required to be included in the review |
| Outcome | Any type of outcome was expected from the included articles regarding the study of the Approach-Avoidance Bias in Virtual Reality |
| Others | No timing restrictions; Virtual Reality and natural setups; Studies reported in English; Published studies in peer-reviewed journals. |

### Information sources

The searches were conducted using Web of Science (WoS; 1945 onwards), PUBMED (1948 onwards), and Scopus (1969 onwards). Additionally, literature search tools such as Research Rabbit (Chandra et al., 2023) and Connected Papers (Eitan et al., 2019) were utilized to enhance the likelihood of identifying all relevant published articles on the topic. Reference lists of the included studies and any relevant reviews were also thoroughly scanned for further sources. Searches were conducted during October 2024.

### Search strategy

Search strategies for systematic reviews were developed using keywords related to Approach-Avoidance Bias, Virtual Reality, and naturalistic setups. The final search strings were refined by reviewing the keywords used in various articles within the field, alongside synonym keywords for Approach-Avoidance Bias, Virtual Reality, and natural or real-world setups. Supplementary Material 1 provides the specific search terms used, along with an example of the search strategy for one database.

### Study records

#### Data management

The literature search results were downloaded from each database and consolidated into an Excel document (Microsoft Corporation) to facilitate collaboration among reviewers during the selection and data extraction phases. After removing duplicates, the refined database was uploaded to ASReview software [34] for screening. A calibration exercise was first conducted to train and refine the screening process before it began. Reviewers were then trained to use the ASReview software before starting the screening.

For data extraction, the authors created a Google Form to gather relevant information from the selected articles. A similar calibration and training procedure was performed before the data extraction phase began.

#### Selection process

The selection process was carried out by three independent reviewers (IRs), under the supervision of two authors (AGC & JMC). Initially, the IRs jointly screened several articles as a calibration exercise. Once a high level of consistency was reached (95%), the IRs proceeded to independently screen the remaining titles and abstracts found in the search, using the inclusion criteria within the ASReview software. The software was trained to maximize the benefits of its machine-learning algorithm. Complete reports were obtained for all titles that met the inclusion criteria (Table 1) or where uncertainty remained. The IRs then reviewed the full text of these articles to determine whether they should be included. Any disagreements were resolved through discussions between the IRs and the principal review authors. All reasons for excluding articles were carefully documented. Both the review authors and the IRs had access to the journal titles, study authors, and institutions involved.

#### Data collection process

The review authors (AGC, JMC, & PK) created a Google Form questionnaire and a detailed instruction manual to guide the extraction of relevant information from each included article. Following a training and calibration phase, three IRs carried out the data extraction under the supervision of AGC and JMC. The Google Form contained specific questions designed to address various aspects of the research question. Any disagreements during the extraction process were resolved through discussions between the IRs and the authors.

### Data items

In this review, various types of information were extracted from all included articles. These included bibliographic details such as author names, journal titles, year of publication, and DOI. We also collected specific information about the study content, including the main objectives and the nature of the research. Additionally, paradigm-related details, such as the type of task and stimuli presented, were recorded. Given the focus of this review, methodological information was also gathered, including the devices employed, the type of action performed, and behavioral data recorded. Finally, we extracted details about the reported limitations and suggestions for future research related to using VR to study the AAB. A comprehensive list of data items can be found in Supplementary Material 2.

### Risk of bias

The JBI Critical Appraisal Tools [35,36] were used to assess the potential risk of bias for each study. For each article, the tool’s corresponding extension regarding the experimental design was employed. Two authors (AGC & JMC) conducted this step.

### Data synthesis

This review contains text, tables, and figures to summarize and explain the primary characteristics and findings of the included studies. The narrative synthesis method allowed us to establish relationships and findings between the included studies [37]. Due to significant heterogeneity among the included studies—in their aims, designs, and methodological characteristics—as well as inconsistencies in measured outcomes, a meta-analysis was not conducted [38,39]. The lack of uniformity across studies would make establishing reliable comparators challenging and could lead to potentially misleading generalizations. Consequently, a narrative synthesis was favored to better capture the nuanced insights each study provided.

## Results

### Study selection

Following the initial search, 413 articles were found to be potentially eligible. The initial number of articles was reduced to 327 after removing duplicates. To select the articles, we conducted two screening processes. The first screening was conducted based on the information presented in the article’s title, and 188 articles were excluded. Then, we conducted the second screening, grounded on the information provided in the article’s abstract, 167 articles were excluded in this stage. The search conducted in other browsers gave results that duplicated the articles found in the other databases; hence, they were not included. Another 7 articles were excluded during the data extraction process since they did not fulfill one or more of the specific criteria for this review. Figure 1 illustrates the number of considered articles through the different stages.

**Figure 1.**
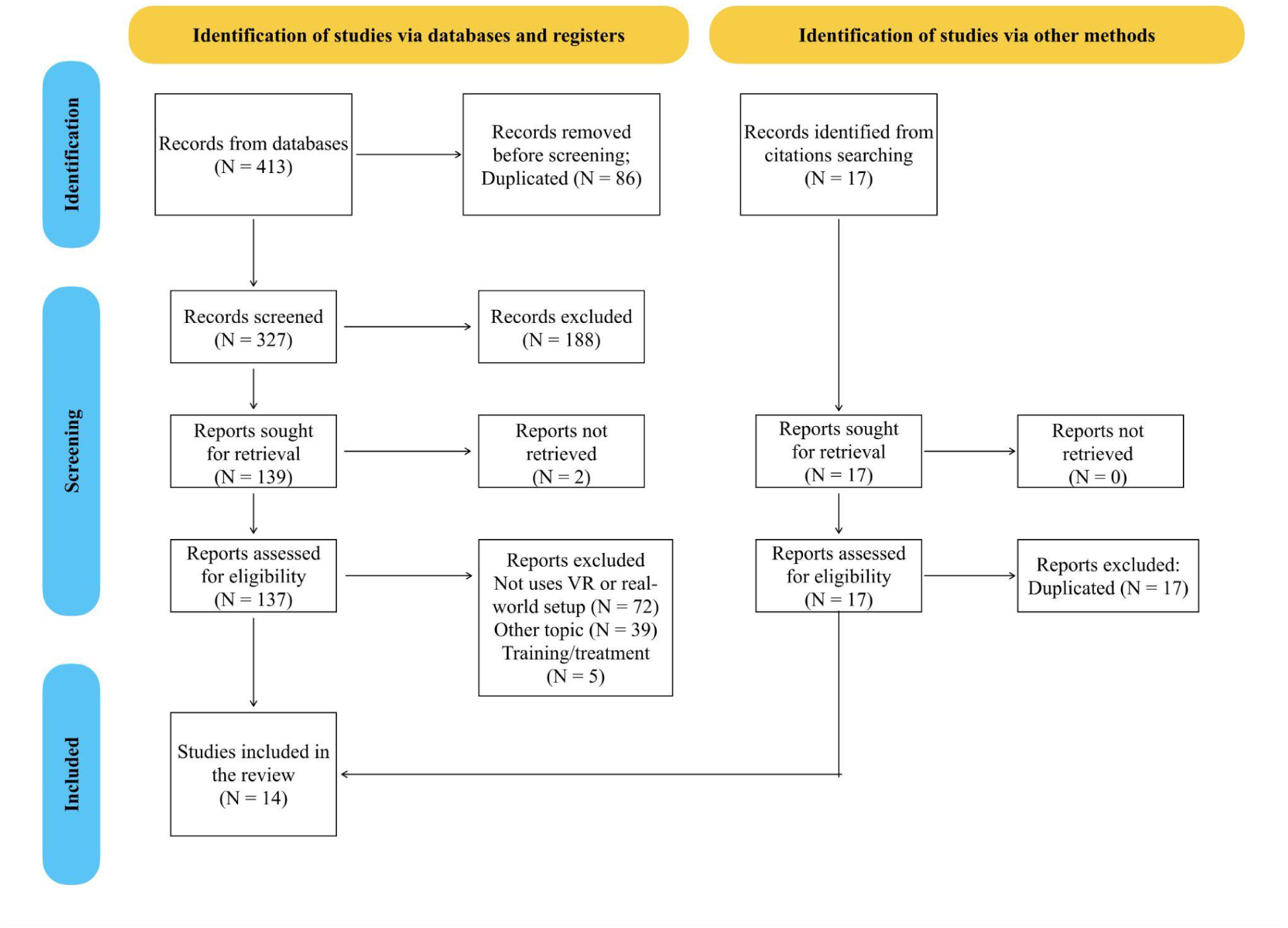
PRISMA flowchart.

One of the main reasons for excluding articles was the absence of the use of VR or natural/real-world setup to study the bias (N = 72). Secondly, many articles (N = 39) were excluded because they were unrelated to the AAB or because they aimed at studying automatic implicit bias (AIB). The AAB refers to the instinctive tendency to move toward positive stimuli and away from negative ones, often guided by perceived emotional valence, like pleasure or threat. In contrast, AIB involves unconscious associations or stereotypes toward groups or categories (e.g., race, gender) that shape perception and behavior without conscious awareness [40–42]. While approach-avoidance bias relates to immediate physical or emotional responses to stimuli, implicit bias operates through ingrained attitudes that subtly influence decisions and judgments. Finally, it is possible to see many studies using VR (N = 44) that were not included since they used the VR environment as a training or treatment. However, this shows the interest in moving towards more realistic setups.

### Studies characteristics

The publication dates of the studies included in this review range from 2016 to 2024. Our results show that an increasing number of articles in the last three years have used VR or a real-world setup to study the AAB (Figure 2).

**Figure 2.**
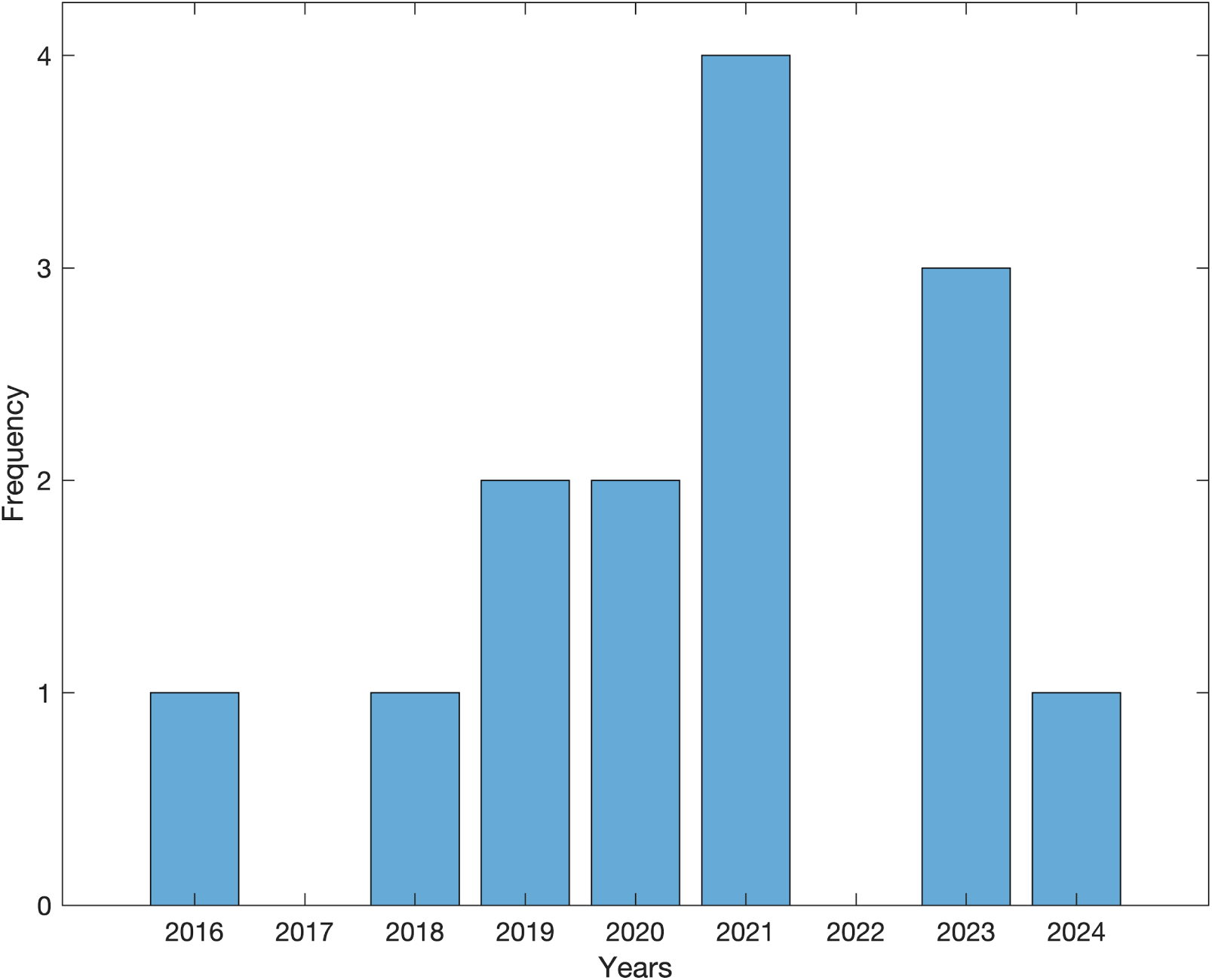
Number of included articles per year.

The included articles are diverse. This is evident since the research question and the main objective of the included studies differ from one to the other. For example, some articles seek to explore behavioral bias towards food items (N = 4), while others explore some domains of social interaction (N = 4). On the other hand, two of the included studies explored differences regarding psychiatric disorders or personality traits. Also, some of the included articles aimed to explore how the Approach-Avoidance Behaviors are modulated when using a VR environment (N = 2). Other studies explored fear (N = 1), gaming disorder (N = 1), and alcohol intake (N = 1). The diversity of the research question and objective of the included studies highlights the versatility of the AAB framework to be applied to different research topics.

The diversity regarding the research question is also translated into a variation, such as the articles included. In this sense, some of the included articles aimed to explore behavioral aspects of the AAB linked to different conditions (e.g., social interactions and food preference, among others), while other articles aimed to prove the feasibility of studying the AAB in VR. However, some articles addressed both behavioral and feasibility aspects simultaneously. These articles, respectively, can be categorized as *Empirical* (N = 14) and *Methodological* (N = 7) regarding their primary objective. All the included articles had a question or objective aiming to explore the bias in VR. At the same time, only half of them were also aimed at testing methodological aspects of moving the study of the AAB from a desktop-based setup to VR.

### Risk of Bias

Table 2 summarizes the risk of bias evaluation outcomes, showing that most included articles meet the tool’s criteria for medium to high quality. Although minor details were occasionally missing from the experimental setup descriptions, this did not impact the studies’ clarity or reproducibility. Additionally, only studies that conducted control measurements of the VR setup (e.g., desktop-based or real-life comparisons) or included comparisons across different population groups were categorized as case-control studies. Detailed explanations for each criterion are provided in Supplementary Material 3.

**Table 2:**
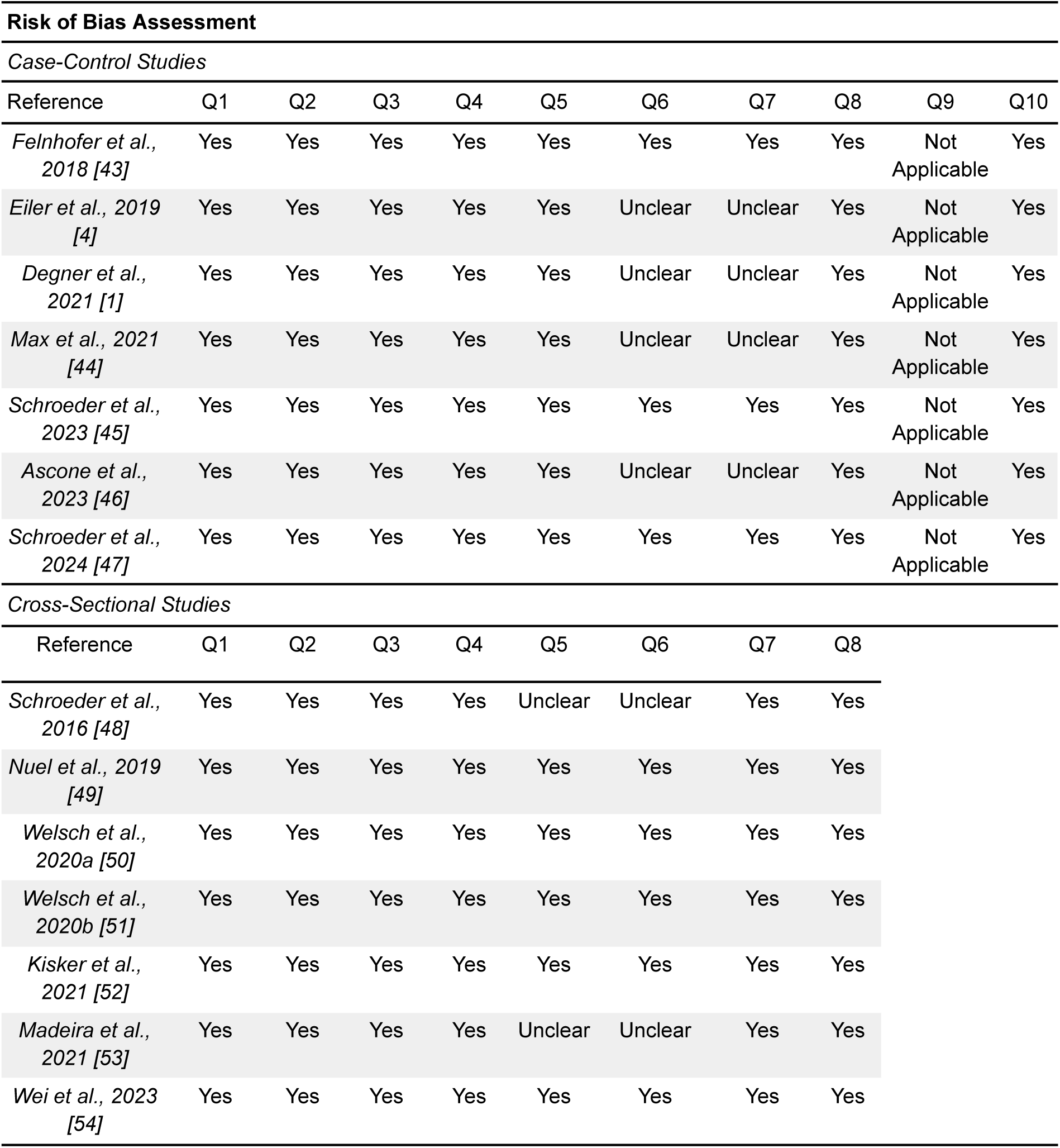

| **Risk of Bias Assessment** |  |  |  |  |  |  |  |  |  |  |
| --- | --- | --- | --- | --- | --- | --- | --- | --- | --- | --- |
| *Case-Control Studies* |  |  |  |  |  |  |  |  |  |  |
| Reference | Q1 | Q2 | Q3 | Q4 | Q5 | Q6 | Q7 | Q8 | Q9 | Q10 |
| *Felinhofer et al., 2018 [43]* | Yes | Yes | Yes | Yes | Yes | Yes | Yes | Yes | Not Applicable | Yes |
| *Eiler et al., 2019 [4]* | Yes | Yes | Yes | Yes | Yes | Unclear | Unclear | Yes | Not Applicable | Yes |
| *Degner et al., 2021 [1]* | Yes | Yes | Yes | Yes | Yes | Unclear | Unclear | Yes | Not Applicable | Yes |
| *Max et al., 2021 [44]* | Yes | Yes | Yes | Yes | Yes | Unclear | Unclear | Yes | Not Applicable | Yes |
| *Schroeder et al., 2023 [45]* | Yes | Yes | Yes | Yes | Yes | Yes | Yes | Yes | Not Applicable | Yes |
| *Ascone et al., 2023 [46]* | Yes | Yes | Yes | Yes | Yes | Unclear | Unclear | Yes | Not Applicable | Yes |
| *Schroeder et al., 2024 [47]* | Yes | Yes | Yes | Yes | Yes | Yes | Yes | Yes | Not Applicable | Yes |
| *Cross-Sectional Studies* |  |  |  |  |  |  |  |  |  |  |
| Reference | Q1 | Q2 | Q3 | Q4 | Q5 | Q6 | Q7 | Q8 |  |  |
| *Schroeder et al., 2016 [48]* | Yes | Yes | Yes | Yes | Unclear | Unclear | Yes | Yes |  |  |
| *Nuel et al., 2019 [49]* | Yes | Yes | Yes | Yes | Yes | Yes | Yes | Yes |  |  |
| *Welsch et al., 2020a [50]* | Yes | Yes | Yes | Yes | Yes | Yes | Yes | Yes |  |  |
| *Welsch et al., 2020b [51]* | Yes | Yes | Yes | Yes | Yes | Yes | Yes | Yes |  |  |
| *Kisker et al., 2021 [52]* | Yes | Yes | Yes | Yes | Yes | Yes | Yes | Yes |  |  |
| *Madeira et al., 2021 [53]* | Yes | Yes | Yes | Yes | Unclear | Unclear | Yes | Yes |  |  |
| *Wei et al., 2023 [54]* | Yes | Yes | Yes | Yes | Yes | Yes | Yes | Yes |  |  |

### Paradigm Characteristics

We reviewed paradigm-related experimental characteristics from each one of the included studies. These characteristics involve the type of task^1^ used to study the bias, the stimuli the participants had to interact with, the bodily response given by the participant, the type of behavioral data collected, and the presence of a comparison condition (in methodological terms). By systematizing these characteristics, we can provide insights into the different approaches used to study the bias in VR.

We found a substantial diversity of stimuli used to measure the AAB across all included articles. Most of the included articles implemented 3D stimuli in their experimental designs, and only one used 2D images within the VR environment. Regardless of the actual content of the stimuli or prime used by each study, which is highly related to its research question, 13 out of the 14 included studies used 3D stimuli displayed in the VR environment. A variation similar to the research question was found in terms of the content since each article incorporated different stimuli considering its primary objective. Examples of the used stimuli are 2D images of alcoholic vs. non-alcoholic beverages, 3D virtual representations of food vs. non-food items, smoke vs. non-smoke items, and virtual avatars. Across studies, stimuli served different contextual purposes, such as mimicking real-world social interactions, ecological tasks, and fears by allowing participants to interact with virtual objects, agents, and environments. Despite this diverse implementation of stimuli, an observed commonality concerns the common goal of evoking context-specific automatic responses relevant to approach and avoidance tendencies.

Also, we found that the tendency to include bodily movements as responses is present in all of the included articles. All the included articles employed some level of embodied movement in the responses from the participants. From pushing/pulling movements (N = 5) using, for example, a joystick, which is one of the most common types of movements used when studying the AAB or a hand gesture, to stepping or freely walking (N = 7), grabbing (N = 5), and even tapping (N = 1) and engaging in verbal conversations (N = 1) (Figure 3). These results show the multiplicity of actions that can be explored beyond pushing and pulling when it comes to studying the embodied mechanisms of the AAB, which encourages thinking about new directions to consider, especially when trying to study the AAB as it occurs outside the laboratory. Similarly, it highlights the versatility of using a VR environment when studying embodied dynamics since the possibilities of movements are more extensive than those implemented in desktop-based experiments.

**Figure 3.**
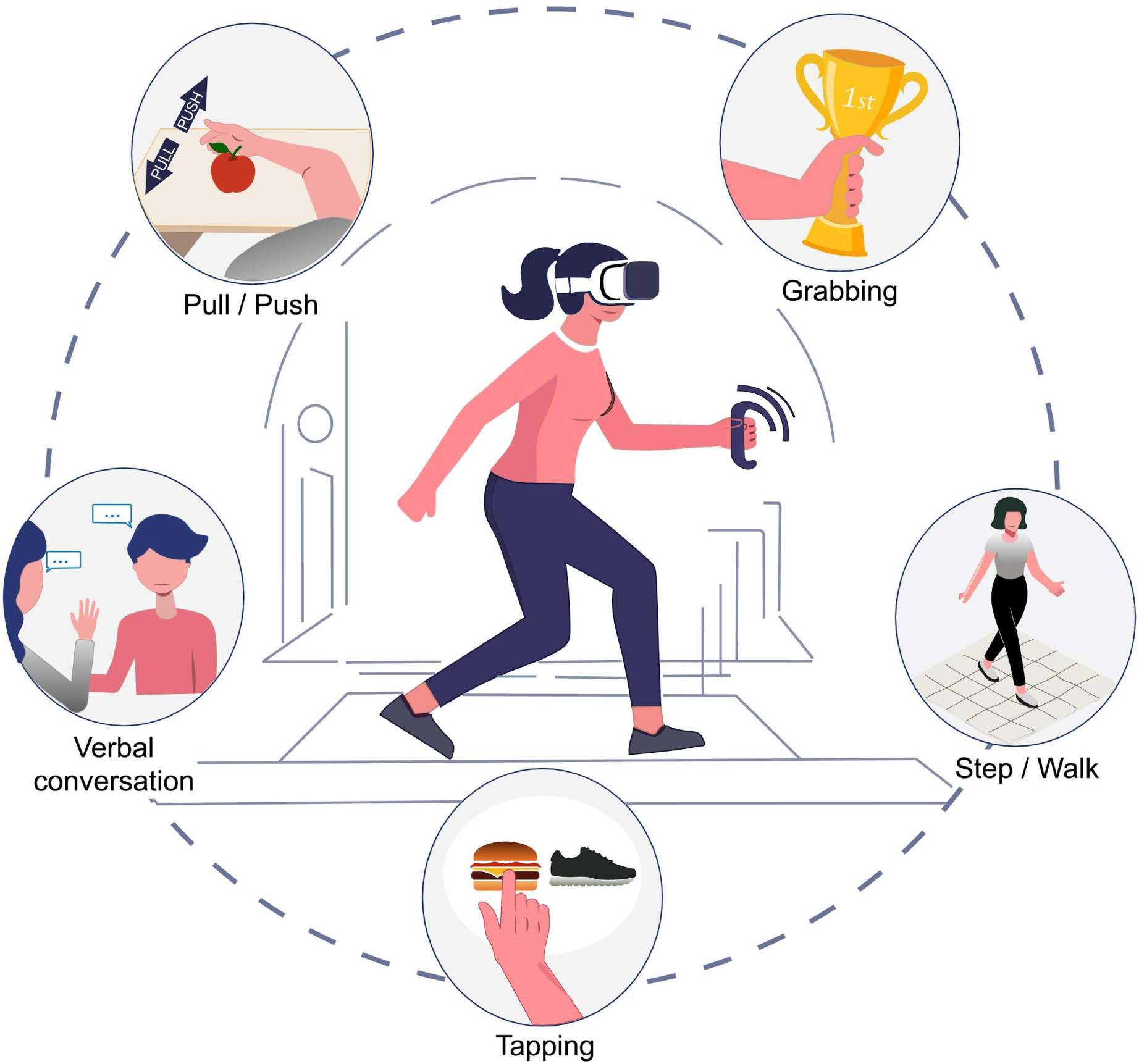
Graphical representation of embodied movements used in virtual reality (VR) to study approach and avoidance bias (AAB). The central panel of the image shows a subject immersed in a VR environment. The five actions around the circle display the different embodied movements used in the included studies to measure participants’ responses.

Regarding the behavioral data collected, reaction time is one of the primary measures for the AAB. Coherent with previous research in desktop-based computers, reaction time was one of the most common behavioral measures used by the VR studies (N = 11). It is important to notice that some experiments recorded more than one behavioral measure, so they performed multiple analyses. The type of action performed by the participants was also a standard measure (N = 4), followed by the position or distance of the participants in the VR environment (N = 3), the exploration time (N = 2), and the velocity (N = 1). This exemplifies that, even when the bias has been studied mainly as a reaction time-based task, different behavioral components can shed light on the understanding of the dynamics of the bias.

Finally, we found that most of the included studies did not incorporate a control task in terms of setup. Of the total reviewed studies, only four conducted a control experiment where the bias was explored in a desktop-based AAT condition to have a control measure or determine differences with the data collected from VR. Three out of the four studies with a control condition used the same stimuli presented in VR for the desktop condition (most of the time, using screenshots of the same objects). This allowed the evaluation of the stimuli as a valid prime to elicitate the bias.

Supplementary Material 4, available through the Open Science Framework (OSF)^2^, provides details about the paradigm characteristics described here.

### Technical Characteristics

Due to the methodological emphasis of the present review, we included the systematization of characteristics related to technical aspects of the experimental setups. We extracted characteristics from the used VR headsets (e.g., brand, sampling rate), controller type, participants’ position and possibility of movement during the experiment, and the system to generate or display the environment. We assessed if any type of (neuro)physiological data was acquired during the experiment (e.g., brain or cardiac activity, skin conductance, among others). The information provided here is based on what is reported in each article; no information was included through research. The systematization of these characteristics might promote and guide the development of new experimental setups. A detailed description of these categories can be found in Supplementary Material 4.

Regarding the type of VR device used for displaying the environment, Head Mounted Displays (HMD) were the most common ones. Only two of the included studies implemented glasses instead of classical HMD VR equipment. Brands and manufacturers varied across the studies, being *Vive* being the most common one (N = 5). The resolution of the environment and the sampling rate for collecting behavioral data (e.g., position) were also diverse for the different studies. However, 7 of the included articles did not report resolution information, and eight did not offer any insights regarding the sampling rate. Most of the controllers were apparatuses from the same brand as the HMD. Six of the systems used for creating and displaying the virtual environment implemented some version of Unity, while two used Vizard. There was no information regarding this parameter in four of the included studies.

Only two of the included studies performed measures concerning the collection of (neuro)physiological activity measuring brain activity. One study implemented fNIRS, while the other used EEG and ECG.

Finally, regarding the physical positioning of the participants during the experiment, it was more common for them to be sitting down and performing a hand-arm movement (N = 6) or standing while stepping or walking (N = 6). In one study, participants performed pulling and pushing movements despite using an experimental design involving a standing position. Similarly, one study required participants physically position to be seated but engage in a verbal conversation. Only one study incorporated both leaning movements (e., forward and backward body inclinations) while participants were seated and stepping movements (e.g., forward and backward steps) while participants maintained a standing position.

### Limitations and Future Direction

This review analyzes studies using VR or real-world paradigms to examine the AAB. To keep this focus, we report only the limitations and future directions from the included articles that directly pertain to the implementation of VR or real-world setups. This review does not include other types of limitations, such as sample size or characteristics, and broader suggestions (e.g., investigating setup feasibility as a treatment or in other populations). Accordingly, Table 3 presents limitations and future directions from the articles that specifically impact the application of VR or real-world setups for studying the AAB and might be helpful for the development of future research.

**Table 3:**
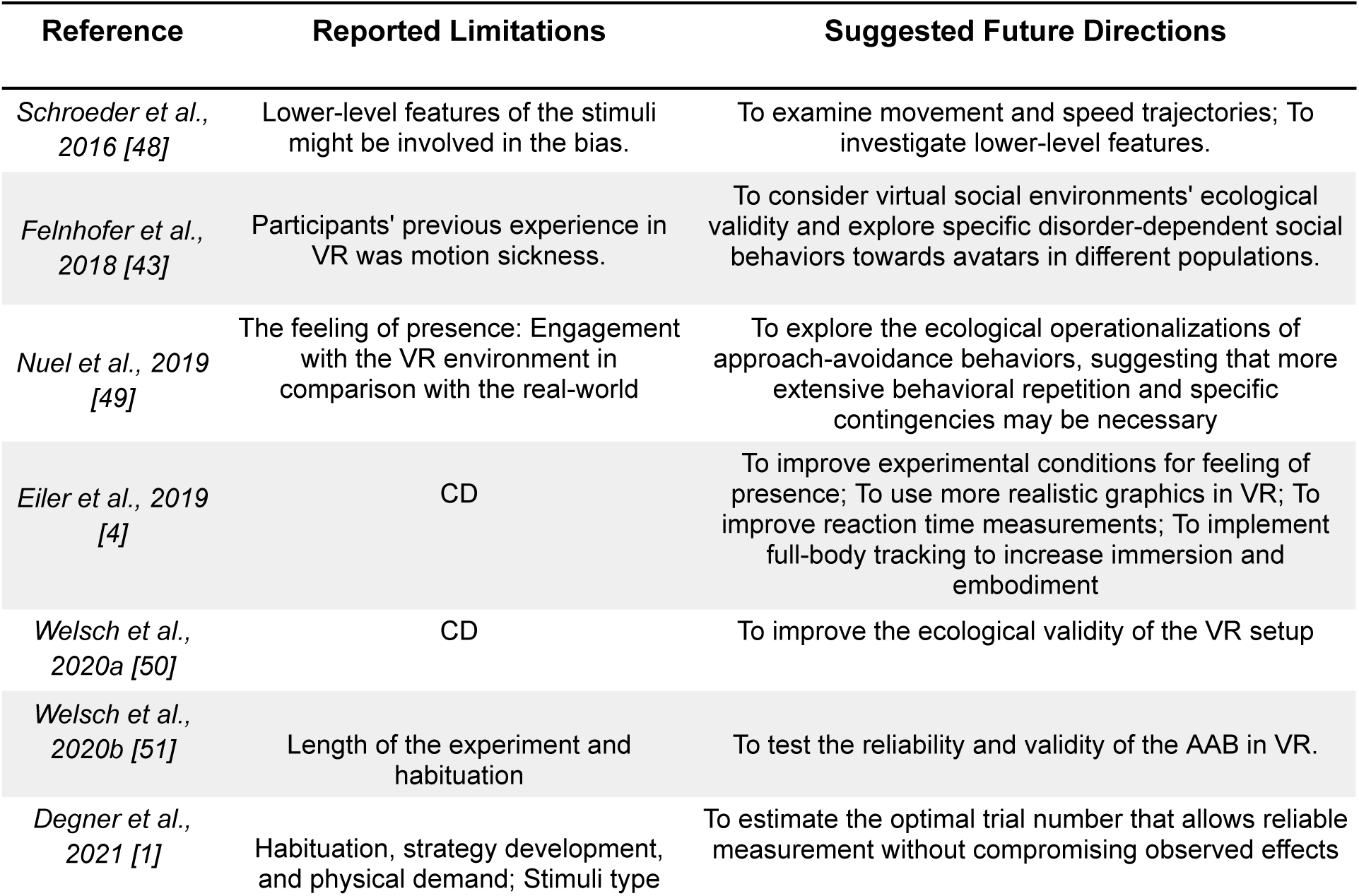
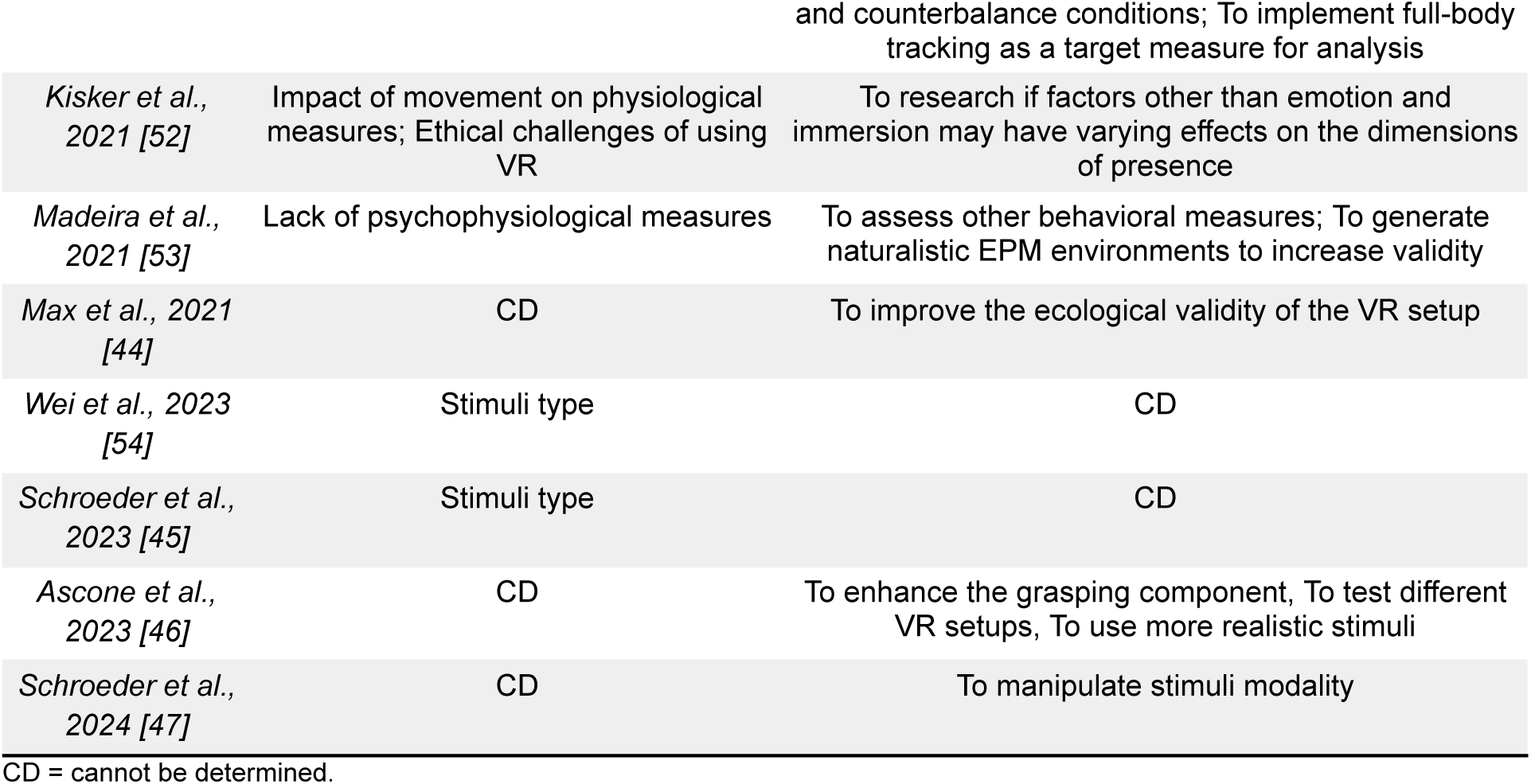

## Discussion

This systematic review provides a detailed overview of the current literature utilizing VR or real-life paradigms to investigate the AAB. By examining the diverse paradigm and methodological characteristics, we offer a narrative synthesis of evidence that supports the feasibility of implementing the AAB in more flexible, ecologically valid environments while preserving its behavioral properties. The synthesized information reveals the significant challenges of transferring the AAB from traditional paradigms, highlighting a wide range of stimuli, embodied responses, and collected behavioral data. The studies included in this review varied regarding virtual environments, devices, resolution, and sampling quality. In the following section, we will discuss additional factors observed during this review and the implications of our findings on the AAB research. We will explore the methodological considerations reported for translating traditional paradigms into virtual environments and examine the impact of diverse stimuli and response measures on research outcomes. We will also highlight potential future research directions, emphasizing the importance of methodological consistency, ecological validity, and additional behavioral measurements in understanding the AAB within immersive settings.

### Divergent studies

While the primary aim of this systematic review was to examine how the study of AAB has been implemented using VR and real-world settings, a substantial portion of the literature diverges toward treatment and training applications. These studies often suggest that the AAB can be effectively transferred into VR without delving into its underlying behavioral components or the suitability of the measurements collected. Examples include Jahn et al. [55] and Smeijers et al. [56], which focus on health-related outcomes and aggressive impulse management. Additionally, many studies explore automatic implicit biases, which often are modulated by other factors like moral judgment rather than directly examining the AAB. For instance, studies by Banakou et al. [57], Kishore et al. [58], Peck et al. [59], You et al. [60], and Persky et al. [61] investigate racial bias through virtual embodiment techniques. Echoing this line of inquiry, Banakou et al. (2020) examined how social context affects the feeling of owning one’s body and contributes to implicit racial bias. This orientation reflects a broader trend, where VR research is primarily used to address not only socio-psychological biases related to ethnicity but also physical disability [62] and social anxiety [63]. While such applications are valuable, they fall outside the immediate scope of understanding the AAB’s behavioral nuances as they occur in real-world dynamic situations. By assuming the methodological and behavioral elements, movements, and technologies involved, these studies neglect the crucial need for a foundational common framework that would eventually allow accurate translation of the AAB research into VR. Without this groundwork, the legitimacy of results and their application to clinical populations remains questionable, as the essential task of establishing baseline principles for VR integration into the AAB research has not been adequately addressed. Together, these divergent studies highlight a gap in the literature where future research should aim to disentangle the behavioral elements that characterize the AAB from broader bias investigations related to it. This approach would ensure that findings directly affect the theoretical understanding and practical deployment of the AAB in VR settings.

### Clinical populations

Besides measuring cognitive biases, improving methods (or developing novel ones) for intervention and treatment of AAB-related biases is crucial for counteracting their negative impact (e.g., substance abuse) on individuals and groups. Effective interventions could help individuals recognize and modify their biases, fostering more balanced behaviors. Some of the studies included in this review investigated AAB tendencies using VR within clinical populations for therapeutic purposes. These studies focused on diverse conditions such as psychopathy [50], alcohol abuse [46], internet gaming disorders [54], and habitual food avoidance in anorexia [47], highlighting variations in inhibitory control mechanisms and demonstrating the feasibility of conducting AAB research using VR setups that extend beyond traditional methods like bottom presses and joystick manipulation. However, our understanding of the bias itself within real and virtual contexts remains limited due to the recent emergence of such studies, which vary significantly in their methodologies and aims. This diversity highlights a gap in the literature, underscoring the need for further research to comprehensively examine the AAB within these environments to better grasp its fundamental characteristics and applications. Further research in this area can contribute to establishing flexible therapeutic interventions and developing preventative strategies that target bias formation and maintenance.

### Mapping the Terrain: A Visual Examination of the Relationships on AAB Research

The benefits of understanding how an AAB occurs in natural dynamic environments are undeniable, particularly as interest in this area grows with state-of-the-art technologies. The ability to simulate realistic settings using VR opens new avenues for exploring the AAB with greater ecological validity; however, this may also lead to less standardized methodological procedures. The interest in novel approaches to the AAB using VR and deviating from traditional paradigms has increased over the years (Figure 2). This shift has led to diverse methodologies and designs that illustrate complex interactions between experimental stimuli used and embodied responses measured. Figure 4 presents an organized overview of the studies in this review, clearly delineating those employing 2D versus 3D stimuli involving sitting or standing environments. Notably, most studies that address empirical and methodological dimensions utilized sophisticated stimuli, including virtual avatars and interactive objects, to elicit a broader range of behavioral responses. Although studies shared the fundamental assumption that changes in physical distance (increase and decrease) modulate the AAB [64], these changes were measured via distinct behavioral metrics. Traditional measures like reaction time were most common among studies addressing only empirical aspects of the AAB within sitting scenarios that facilitated the physical interactions with the virtual stimuli. Mixed and distance-related measurements collected as behavioral responses emphasized the influence of both real and virtual settings on participant behavior and were most often used within standing scenarios. Notably, most studies that used alternative approaches (e.g., standing scenarios, avatar interaction, and mixed or distance-related responses) aligned toward more coherent experimental designs, prioritizing the ecological niche of the behavioral element involved. The conceptual map illustrates a clear distinction between coherent and incoherent paradigms, indicating that while some studies successfully aligned participants’ embodied responses in VR with their physical, real-world position (such as matching stepping actions in the VR environment with actual stepping while standing) and employed devices like motion-tracking cameras that preserved the integrity of the movement, other studies failed to maintain this coherency. This relational map highlights the necessity for future research to ensure coherent experimental setups to enhance the findings’ validity and applicability. In the future, it will be essential to prioritize methodological rigor when developing VR environments to achieve consistent and reliable measurements of the AAB, fostering more nuanced insights into the complex behavioral characteristics of the AAB.

**Figure 4.**
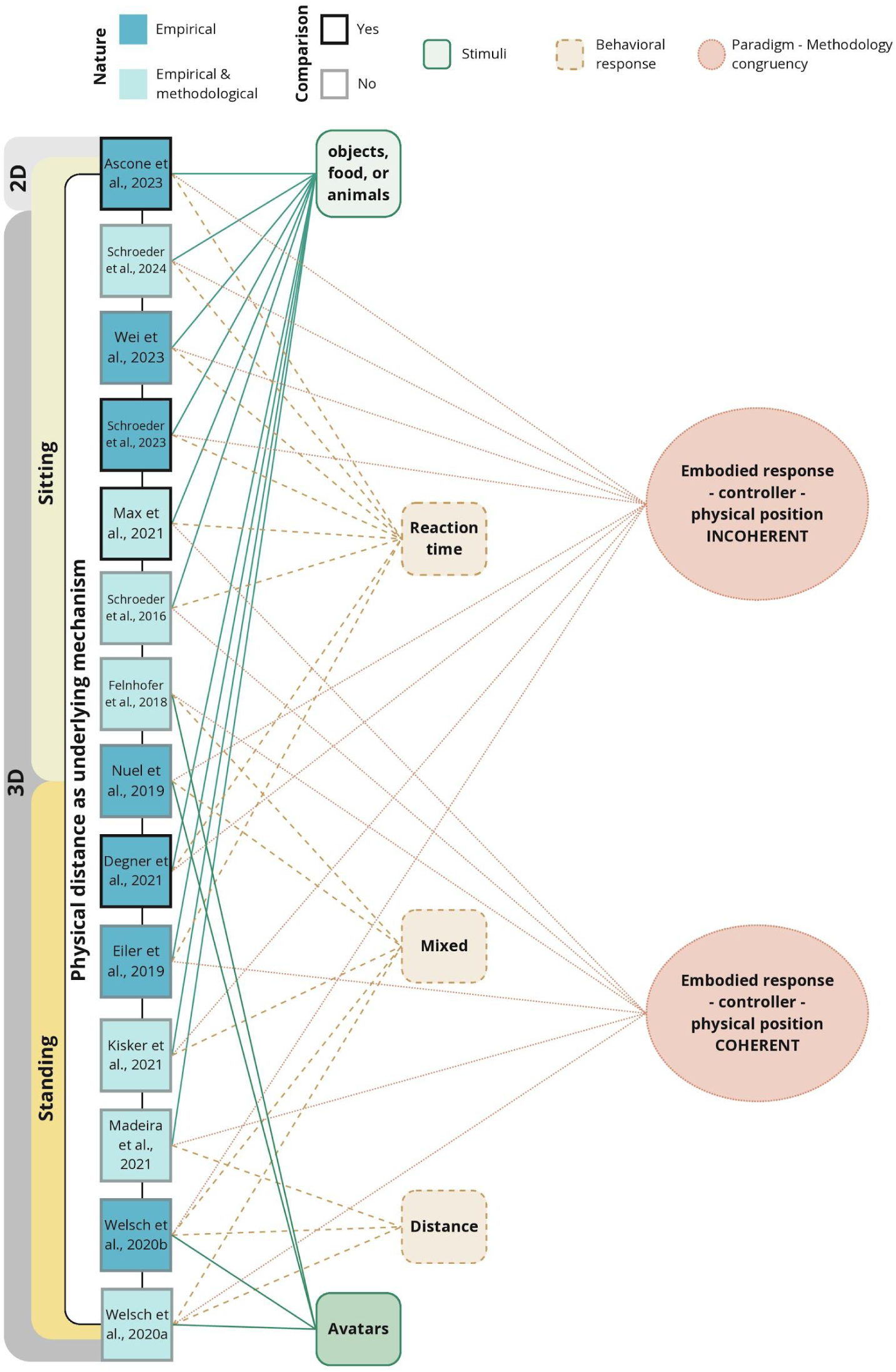
A conceptual map illustrating the relationship between all included articles (labeled by author and year), the types of stimuli (green), and the behavioral responses (orange) measured within the approach and avoidance bias research. From left to right, the map characterizes studies into empirical and methodological, distinguishing between 2D and 3D stimuli settings. It contrasts those studies that used small stimuli (such as objects, food, and animals) with those that promoted whole-body interaction with virtual avatars and associated types of behavioral responses measured. Paradigm-methodology congruence identifies whether each study’s embodied responses and devices used to mimic movements in virtual reality were coherent or incoherent with participants’ physical position. Lines denote the interactions of each study with the categories.

A key finding derived from this relational examination of the different approaches to studying AAB using VR concerns the importance of coherent design paradigms and their methodological implementations. Eiler et al. [4] emphasize the need for stimuli with distinctive yet subtle features, allowing them to remain recognizable despite rapid automatic actions. Coherence between stimuli and participants’ physical movements, such as grasping and stepping, must be maintained to ensure meaningful interactions within the 3D space. For instance, if the AAB is assessed by measuring stepping behavior in VR, participants should be physically standing, with actual stepping movements captured, rather than using controller inputs. Similarly, grasping actions should be facilitated by actual hand movements rather than button presses. Solzbacher et al. [15] demonstrate that designs promoting such coherent interactions of, for example, pulling and pushing (using a joystick vs. a response pad) show a significant reaction time advantage, underscoring the critical bodily component of the AAB. Moreover, as Keshava et al. [65] show, realistic action affordances impact goal-oriented planning, meaning that experiments should preserve the integrity of physical and virtual movement affordances. Thus, experimental designs integrating coherent action simulation can provide deeper insights into the AAB’s underlying mechanisms and improve the effectiveness of therapeutic treatment.

However, this dichotomy between experimental design and afforded physical actions still remains implicit to current AAB research. For instance, some bodily responses used to measure AAB such as engaging in verbal conversations [43] and tapping [47] deviate from the idea of an embodied component inherent to AAB that is rooted in its evolutionary role [6,15]. These movements seem disconnected from the actual AAB tendencies they aim to assess and are unlikely to activate the core behavioral mechanisms underlying the AAB. This misalignment suggests that tasks such as conversing and tapping may not effectively engage the intuitive and automatic action-oriented responses that AAB is theorized to evoke. Instead, these tasks risk focusing on cognitive processing rather than eliciting genuine AAB tendencies. In Cognitive-Bias Modification (CBM), Van Dessel et al.’s [66] ABC training simulates real-life situations to aid individuals in making goal-relevant behavioral choices (to reduce alcohol addiction) in response to antecedent cues and their action consequences. However, joystick-based pulling and pushing in ABC training does not accurately reflect real-life behaviors associated with the approach and avoidance of alcohol. The incoherence between the ABC training method and real-life behavior may lead to participants experiencing cognitive dissonance. If individuals are trained to push and pull images rather than engage in actions that mimic actual drinking behaviors, they may struggle to internalize the training as they may not develop the necessary skills to handle actual situations involving alcohol consumption. Similarly, Markmann and Brendl’s [67] interpretation of the manikin task highlights how incoherent movements in experimental designs can obscure a proper understanding of AAB. They show a compatibility effect that relies on participants’ spatial representation of themselves rather than on their physical actions. This misalignment between conceptual and physical actions may lead to results that do not genuinely reflect AAB’s underlying mechanisms, emphasizing the need to align perceptual and motor tasks with participants’ spatial self-representations to maintain the integrity of action-relevant movements. Yet, using pushing and pulling movements to positive and negative words, which are not actions typically associated with such stimuli in real life, questions the reliability of the findings. Therefore, coherent experimental designs that preserve the integrity of physical and simulated movements would advance our understanding of the AAB in everyday life scenarios and, when possible, improve the reliability of physiological data collection.

### The AAB beyond the distance mechanism

Our results align with the standard definition of the AAB as a distance-regulation mechanism. This implies that changes in the distance between an agent and a target object (i.e., increasing or decreasing) account for the behavioral phenomena underlying the bias [10,15,64]. Accordingly, pushing and pulling actions, often conceptualized as arm extension and flexion, are among the most frequently studied behaviors in tasks exploring the AAB in both desktop-based and VR setups [4,15,46,50,54]. However, framing pushing and pulling behaviors within the context of the AAB is not without criticism. One primary concern is the lack of a single, inherent meaning for these actions to be exclusively categorized as approach or avoidance behaviors [68–70]. For instance, retracting the hand toward the body (a pulling movement typically associated with approach) could instead signify avoidance, such as when withdrawing the hand from a heat source [49]. This is also possible when using walking towards (i.e., approach) or away (i.e., avoid) from an object as the behavioral action to study the bias. When only considering the reduction or increase of the distance, the agency of the subject is neglected to a secondary degree. For example, an agent might reduce the distance (considered as an approach behavior) between its own body and an insect (e.g., a spider) to capture or terminate it (an avoidance behavior). These examples illustrate how an action traditionally conceptualized as approach behavior (e.g., arm flexion, walking towards an object) can, in specific contexts, function as avoidance behavior [49]. This underscores the fact that the distance-regulation mechanism is highly context- and stimulus-dependent, a factor that must be considered when investigating the AAB.

On the other hand, these movements are limited in scope, as not all behaviors towards objects can simply be explained by the dichotomies of being pushed, pulled, or approached/avoided through walking towards or away. As Nuel and colleagues [49] argue, a comprehensive conceptualization of the AAB must account for the situated nature of the bias, recognizing the importance of the agent’s history of interactions and its available action possibilities. In this context, the agency of the organism and the affordances of the objects and environment (i.e., possibilities for interaction [71]) should be considered when studying the bias [72–74]. For example, this could involve expanding the range and variety of actions an agent can take when encountering a particular object. In this regard, VR offers researchers a unique opportunity to explore a range of simultaneous interaction options, moving beyond the traditional dichotomy, by observing the agent’s natural behavior as they interact with the presented stimuli.

### (Neuro)physiology of the AAB in VR

The systematization of the literature shows a need for the inclusion of (neuro)physiological measurements when studying the AAB in VR. Of the 14 reviewed articles, only two [44,52] incorporated physiological measurements, focusing on brain activity. This underscores a need for further exploration of (neuro)physiological dynamics in more ecologically valid settings related to the AAB. Notably, while studies on the (neuro)physiological dynamics of the AAB are relatively scarce, most studies exploring the (neuro)physiological dynamics are restricted to computer-based setups. Furthermore, stationary neuroimaging techniques are often used, with a clear tendency for systems that provide high spatial resolution data compared to others (e.g., functional Magnetic Resonance Imaging; fMRI) [3,75,76]. This tendency aligns with the standard analysis previously conducted on the AAB, which is mainly related to activity in the amygdala due to its involvement in motivation and emotional responses [77–79]. The limited application of (neuro)physiological measurements in VR settings may stem from the challenges of integrating high spatial resolution neuroimaging technologies with immersive VR setups [80]. This presents an opportunity for further research to address these technological and methodological barriers.

The AAB has also been explored using other neuroimaging techniques. Frontal Alpha Asymmetry (FAA) is one of the most commonly implemented analyses when studying the AAB using Electroencephalography (EEG) [81,82]. FAA has been primarily related to emotion, motivation, and cognitive control [83–85], which are the most common underlying cognitive processes associated with the AAB [70,86,87]. This line of research shows the feasibility of studying the AAB using techniques with lower spatial resolution and higher temporal resolution. Furthermore, it opens the possibility of studying the AAB using technical-methodological frameworks such as Mobile Brain/Body Imaging (MoBI; [88–90]), which allows the study of brain dynamics during mobile tasks involving more naturalistic approach-avoidance scenarios. Hence, studying the AAB in more naturalistic conditions using VR or in real-life setups, posits a complementary approach to classical laboratory setups, allowing the development of tasks in real-world contexts (e.g., simulated or real situations) where individuals have higher freedom for self-agency and interaction [91–97]. This approach will allow the exploration of common EEG markers associated with the AAB, such as FAA, in a more naturalistic and meaningful way as the bias occurs in real life.

### Limitations and Future Directions

The reported challenges and limitations of the included studies are relatively underexplored. Most of the included articles report participant-specific limitations related to using VR for experiments (i.e., ethical aspects, habituation, motion sickness, among others) and limitations regarding stimulus type and its characteristics. However, these discussions often overlook the pivotal technical and methodological considerations for transferring the AAB into more naturalistic setups. For example, Eiler et al. [27] provide a thorough analysis of critical aspects to consider for such translation, including high costs, the need for technological expertise, equipment constraints, and challenges in terms of data accuracy. A comprehensive examination of not only conceptual but also technical and methodological challenges is needed to foster reproducibility across multiple populations and research settings. Furthermore, this can encourage the development of innovative approaches that advance the field. Additionally, given the diversity of topics addressed when studying the AAB in VR, establishing a standardized methodological framework could create consistency across studies and enable researchers to build upon one another’s work.

Finally, considering future directions for studying the AAB in VR, many included studies emphasize the need to address embodied aspects of virtual environments to replicate reality better. Also, integrating (neuro)physiological measurement could provide valuable insights into the neural and motor processes underlying the AAB. This approach would allow researchers to explore its (neuro)physiological dynamics in naturalistic interactions, bridging the gap between experimental findings and real-world applications. Furthermore, exploring the AAB in real-life scenarios by placing participants in novel and coherent setups could enhance ecological validity and offer a more nuanced understanding of how VR findings translate to everyday behaviors. Balancing such advancements with scientific rigor and methodological novelty will ensure that research remains robust and reproducible while pushing the boundaries of what can be achieved in this field.

### Conclusions and General Recommendations for Translating the AAB into VR

The results from the present review show a strong tendency to translate the AAB into more naturalistic setups, whether to study the bias itself or for applications such as training or therapeutic interventions. However, when it comes to understanding the bias, the field remains diverse in its research questions and the methodological approaches employed. The following sections will present general recommendations for consistently translating the AAB into VR.

Several of the desktop-based AATs use stimuli that come from databases that have been standardized. Examples of this are the use of pictures from the International Affective Picture System (IAPS; [98]), the Nencki Affective Picture System (NAPS; [99]), or the Image Stimuli for Emotion Elicitation (ISEE; [100]), among others. Using stimuli from these datasets ensures that they have been rigorously tested and validated for eliciting specific emotional states (e.g., positive or negative affect). However, as experimental setups transition to more naturalistic environments, 3D stimuli are often preferred because they provide a more immersive and realistic experience, closely mirroring real-world conditions. Nevertheless, the adequacy of the stimuli to generate the phenomenon that we aim to find, such as the AAB, must remain a critical consideration [101]. To address this, we propose two complementary strategies. First, a desktop-based control condition will be implemented to prove the adequacy of the stimuli in generating the AAB. Some of the reviewed studies that included a control condition (e.g., desktop-based) used screenshots from the VR stimuli in a classical AAT to assess the adequacy of the stimuli without any potential cofounders generated by the VR setup. Moreover, this approach facilitates a systematic examination of the bias, transitioning from controlled experimental setups to more complex, naturalistic environments that closely simulate real-world scenarios [102]. This progression enhances the findings’ ecological validity, relevance, and applicability, bridging the gap between laboratory research and practical, real-world applications. Secondly, it is recommended that validated assessment tools be employed to evaluate key dimensions of the stimuli before their implementation in VR. An example is using the Self-Assessment Mannequin (SAM; [103–105], which can measure dimensions such as pleasure, arousal, and dominance. These measures can help rule out potential confounding factors and ensure the stimuli’s characteristics align with experimental goals. Combining these approaches can enhance confidence in the stimuli’s efficacy while maintaining the rigor of the research, even when the specific stimuli do not belong to a standardized database.

Another essential consideration when designing VR experiments is the type of gesture implemented. Even when a virtual environment already provides a more naturalistic setup, in terms of the stimuli presented, achieving a closer approximation to real-life phenomena requires careful attention to the range of actions participants can perform. Research highlights that the type of embodied response generated significantly influences the phenomena under study [15,65]. Therefore, incorporating coherent interaction between the actions the subject performs inside and outside the VR setup is crucial. For example, suppose that inside the VR environment, the participant has the experience of getting closer to or further away from a target stimulus or avatar by walking. In that case, we encourage the design of experiments where the participants are actually able to perform these movements outside of the environment (e.g., physically stepping forward and backward) instead of using a joystick to control the movement. By incorporating coherent embodied actions that enhance ecological validity and do not artificially restrict the agent’s movements in their interaction with the stimuli, we can assess how affordances for action impact the AAB, providing deeper insights into the underlying mechanisms.

In order to achieve this goal, technical and methodological aspects must be assessed. First, it is important to consider what type of devices are used to record behavioral movements. Following the findings of our review, we recommend prioritizing motion sensors over traditional input devices, like joysticks or response pads, to enhance naturalistic actions since they present fewer restrictions for actions [106,107]. Secondly, graphical features of the virtual environment are equally critical as they contribute to the immersive and embodied experience. For instance, in scenarios involving stepping forward and backward, the environment should dynamically adjust accordingly to the performed movement in order to simulate as much as possible the real-world experience (e.g., realistic object scaling, perspective shifts, detail levels). Similarly, for grasping behaviors, the design should account for interactive elements such as manipulating virtual objects, ensuring that the experience mirrors real-world tactile interactions. By addressing these technical and methodological aspects, we can enhance the implementation of naturalistic actions, leading to a better understanding of the embodied mechanisms of the AAB in the real world.

Finally, we strongly advocate for the collection of multimodal data to enhance the understanding of the underlying mechanisms of biases. Although only two reviewed articles include some type of (neuro)physiological data, its incorporation might provide insights into the foundational processes driving the AAB. For example, it would be possible to explore and understand the role of autonomic physiological states in the decision-making process [19], stress [108], and other dynamics. As we stated before, we recommend formulating research questions targeting this type of data since it will significantly contribute to understanding the basis mechanism of the bias. In this sense, we encourage the development of innovative research questions involving (neuro)physiological dynamics underlying the AAB. Such an approach will facilitate the inclusion of multimodal data such as eye tracking, EEG, and even other measures such as galvanic skin response (GSR) and ECG for a comprehensive understanding of AAB.

## Supplementary Materials

Supplementary Material 1. String of search for Web of Science.

Supplementary Material 2. Data items for data extraction.

Supplementary Material 3. Criteria for quality and risk of bias analysis.

## Authors contribution

Conceptualization, AGC, JMC. and PK; Methodology, AGC; Formal Analysis, AGC; Investigation, AGC; Data Curation, AGC; Writing – Original Draft Preparation, AGC and JMC; Writing – Review & Editing, AGC, JMC, SW, and PK; Visualization, AGC and JMC; Supervision, PK; Project Administration, AGC and JMC; Funding Acquisition, PK and SW.

## Funding

The present systematic review is supported by the University of Osnabrück in cooperation with the Deutsche Forschungsgemeinschaft (DFG, German Research Foundation) in the context of funding the Research Training Group “Situated Cognition” under the project number GRK 2185/2.

## Institutional Review Board Statement

Not applicable.

## Informed Consent Statement

Not applicable.

## Data Availability

Data is available through Open Science Framework: https://osf.io/huv6x/

## Acknowledgments

The authors would like to extend their sincere gratitude to everyone who contributed to the success of this project. They are especially thankful to Franziska Tietze, Niloofar Nazari, and Priyanka Veerabhadrappa for their support in conducting the systematic searches and the screening processes. Additionally, the authors thank Debora Nolte and Tracy Sánchez Pacheco for their insightful feedback and constructive discussions.

## Conflict of interest

The authors declare no conflict of interest.

## Abbreviations List

The following abbreviations are used in this manuscript:

2D: Two-dimensional
3D: Three-dimensional
AAB: Approach and Avoidance Bias
AAT: Approach-Avoidance Task
AIB: Automatic Implicit Bias
CBM: Cognitive-Bias Modification
ECG: Electrocardiogram
EEG: Electroencephalogram
FAA: Frontal Alpha Asymmetry
fNIRS: Functional Near Infrared Spectroscopy
GSR: Galvanic Skin Response
HMD: Head-Mounted Display
IAPS: International Affective Picture System
ISEE: Image Stimuli for Emotion Elicitation
JBI: Joanna Briggs Institute
MoBI: Mobile Brain/Body Imaging
NAPS: Nencki Affective Picture System
PRISMA: Preferred Reporting Items for Systematic Reviews and Meta-analyses
SAM: Self-Assessment Mannequin
SRC: Stimulus-Response Compatibility
VR: Virtual Reality

## Footnotes

1 Due to the large differences between tasks, a systematization is not possible. Hence, we have opted for presenting the description of each task in detail in Supplementary Material 4.

2 The extended data from the results is available in this repository https://osf.io/huv6x/

## Notes

https://osf.io/huv6x/

### Competing Interest Statement

The authors have declared no competing interest.

