## Supplementary Material 1 for "Approach-Avoidance Bias in Virtual and Real-World Simulations: Insights from a Systematic Review of Experimental Setups"

String of search for Web of Science.
((ALL=("approach avoidance bias" OR "approach bias" OR "avoidance bias" OR "motivational bias" OR "approach-avoidance conflict")) OR ALL=("automatic approach bias" OR "automatic bias" OR "approach tendencies" OR "avoidance tendencies")) AND ALL=("virtual reality" OR "VR" OR "immersive environment" OR "virtual environment" OR "augmented reality" OR "mixed reality" OR ‘’natural setup’’ OR ‘’real-world’’).
