## Supplementary Material 2 for "Approach-Avoidance Bias in Virtual and Real-World Simulations: Insights from a Systematic Review of Experimental Setups"

Data items used to extract information from articles:

1. Aim of the study.
2. Nature of the study.
3. Type of AAT.
4. Stimuli used.
5. Embodied response.
6. Behavioral measure.
7. Comparison.
8. VR system.
9. Resolution.
10. Sampling rate.
11. Controllers used.
12. System to generate the environment.
13. Collection of physiological data (in VR).
14. Participant’s position and movement.
