## Supplementary Material 3 for "Approach-Avoidance Bias in Virtual and Real-World Simulations: Insights from a Systematic Review of Experimental Setups"

Criteria for Cross Sectional Studies from the Critical Appraisal tools for use in JBI Systematic Reviews

1. Were the criteria for inclusion in the sample clearly defined?
2. Were the study subjects and the setting described in detail?
3. Was the exposure measured validly and reliably?
4. Were objective, standard criteria used for measurement of the condition?
5. Were confounding factors identified?
6. Were strategies to deal with confounding factors stated?
7. Were the outcomes measured validly and reliably?
8. Was appropriate statistical analysis used?

Criteria for Case-Control Studies from the Critical Appraisal tools for use in JBI Systematic Reviews

1. Were the groups comparable other than the presence of disease in cases or the absence of disease in controls?
2. Were cases and controls matched appropriately?
3. Were the same criteria used for the identification of cases and controls?
4. Was exposure measured in a standard, valid, and reliable way?
5. Was exposure measured in the same way for cases and controls?
6. Were confounding factors identified?
7. Were strategies to deal with confounding factors stated?
8. Were outcomes assessed in a standard, valid, and reliable way for cases and controls?
9. Was the exposure period of interest long enough to be meaningful?
10. Was appropriate statistical analysis used?
